# MacaqueNet: big-team research into the biological drivers of social relationships

**DOI:** 10.1101/2023.09.07.552971

**Authors:** Delphine De Moor, MacaqueNet, Macaela Skelton, Oliver Schülke, Julia Ostner, Christof Neumann, Julie Duboscq, Lauren J. N. Brent

## Abstract

1. For many animals, social relationships are a key determinant of fitness. However, major gaps remain in our understanding of the adaptive function, ontogeny, evolution, and mechanistic underpinnings of social relationships. There is a vast and ever-accumulating amount of social behavioural data on individually recognised animals, an incredible resource to shed light onto the biological basis of social relationships. Yet, the full potential of such data lies in comparative research across taxa with distinct life histories and ecologies. Substantial challenges impede systematic comparisons, one of which is the lack of persistent, accessible, and standardised databases.
2. Here, we advocate for the creation of big-team collaborations and comparative databases to unlock the wealth of behavioural data for research on social relationships by introducing MacaqueNet (https://macaquenet.github.io/).
3. As a global collaboration of over 100 researchers, the MacaqueNet database encompasses data from 1981 to the present on 14 species and is the first publicly searchable and standardised database on affiliative and agonistic animal social networks. With substantial inter-specific variation in ecology and social structure and the first published record on macaque behaviour dating back to 1956, macaque research has already contributed to answering fundamental questions on the biological bases and evolution of social relationships. Building on these strong foundations, we believe that MacaqueNet can further promote collaborative and comparative research on social behaviour.
4. We believe that big-team approaches to building standardised databases, bringing together data contributors and researchers, will aid much-needed large-scale comparative research in behavioural ecology and beyond. We describe the establishment of MacaqueNet, from starting a large-scale collective to the creation of a cross-species collaborative database and the implementation of data entry and retrieval protocols. As such, we hope to provide a functional example for future endeavours of large-scale collaborative research into the biology of social behaviour.

## Introduction

Social relationships are a key predictor of fitness in group-living animals (Hinde, 1976; Snyder-Mackler et al., 2020) and social isolation is an important risk factor for human health and well-being (Holt-Lunstad & Steptoe, 2022). Yet, major gaps remain in our understanding of the adaptive function, evolutionary history, ontogeny, and mechanistic underpinnings of social relationships. Current outstanding questions include which types of social relationships are selected for, and why (Kappeler, 2019; Ostner & Schülke, 2018); what neural and genetic mechanisms underlie social behaviour (Traniello & Robinson, 2021); how social relationships ‘get under the skin’ to impact health, ageing, and survival (Simons et al., 2020; Sarkar et al., 2020) and how these relationships develop over an individual’s lifetime (Siracusa, Higham, et al., 2022) and in the face of adversity, including changing environments in the Anthropocene (Blumstein et al., 2022). While advances are being made to address each of these questions, a holistic understanding of the biology of social relationships will emerge only through the integration of findings across taxonomic groups (NESCent Working Group on Integrative Models of Vertebrate Sociality: Evolution et al., 2014).

Comparative studies are fundamental in understanding the biological basis of traits, as they can reveal broad patterns on evolutionary and developmental history, past and current selective pressures, and underlying mechanisms (Nunn, 2011; Tinbergen, 1963). Yet, a major limitation to comparative research is reaching a sufficiently large and diverse sample for reliable inference (Borries et al., 2016; Schneider et al., 2019). Comparative studies therefore typically rely on combining independent research on individual populations (Lukas & Clutton-Brock, 2017).

There is a vast and ever-accumulating amount of social behavioural data on individually recognised animals (Sheldon et al., 2022). This is an incredible resource for comparative research into the biological basis of social relationships, but it also comes with substantial challenges. The field of animal behaviour historically and currently still consists of many independent research groups collecting and managing data using similar yet slightly different methods (the ‘long tail’ of data: many small datasets that together represent the vast majority of existing data; Wallis et al., 2013). Choices regarding which behaviours to observe, how those behaviours are defined, and which methods are used to record them, result in a wide variation in behavioural data types and definitions. This significantly complicates the creation of standardised databases on social behaviour, compared to existing databases pooling relatively consistently defined and quantified data such as life history, morphological and ecological traits, population sizes and geographic ranges, and measures of biodiversity (SPI-Birds: Culina et al., 2021; The Global Biodiversity Information Facility: Edwards et al., 2000; panTHERIA: Jones et al., 2009; The Primate Life History Database (PLHD): Strier et al., 2010).

Moreover, most comparative studies in animal behaviour typically consist of one-off comparative research projects, spearheaded by an individual research team, and focused on a specific question. The generated databases are usually not conceived for general use, making them difficult to find or repurpose for future studies (O’Dea et al., 2021). As such, the painstaking effort of searching the literature for suitable data, acquiring such data, and standardising the data and metadata across datasets needs to be repeated for each new comparative study (Poisot et al., 2019). A far more sustainable approach is to create enduring and reusable databases, with well-defined protocols for (meta)data standardisation, archiving and accession (Culina et al., 2021). Clear and concise guidelines for such databases are provided by the FAIR guiding principles to ensure data are Findable, Accessible, Interoperable and Reusable (‘FAIR’, Wilkinson et al., 2016). The development of such databases has become a priority for many scientific fields and has led to the creation of big-team science initiatives, bringing together scientists “across labs, institutions, disciplines, cultures, and continents” (Forscher et al., 2022, p. 2). The aim of these collaborations is to enable large teams of researchers to collaborate more effectively and to accelerate scientific discoveries by making it easier to integrate and compare data from different sources.

Many of these endeavours have taken the approach to standardise data collection across research sites and groups, following the ‘Many Labs’ approach (Klein et al., 2014; e.g. ManyPrimates: ManyPrimates et al., 2019; ManyBabies: Visser et al., 2022). This is a promising avenue for future research, and one that ideally should be adopted more broadly at long-term research sites to allow for the identification of broad patterns across sites and study systems (Rubenstein & Abbot, 2017). Yet, it would be a waste (Purgar et al., 2022) to stop using the 70+ years-worth of social behavioural data that already exist (Sheldon et al., 2022). Although complex, there is great potential in standardising existing data and making them usable for comparative research.

The Animal Social Network Repository (ASNR) was the first significant effort to consolidate social behaviour data across different animal species using FAIR principles (Sah et al., 2019). Since its publication, the ASNR database has been used for several comparative studies across a broad taxonomic range (e.g. Collier et al., 2022 and Gagliardi et al., 2023), demonstrating the value of such databases for the scientific community. However, the ASNR database’s significant diversity also presents challenges for comparative research as it contains social data collected in various ways, with substantial variation in observation effort and capturing fundamentally distinct aspects of sociality. A second, recently developed database is DomArchive, which focuses on dominance interactions within groups of various taxa (Strauss et al., 2022). Social dominance is conceptually similar and measured fairly consistently across taxa (Strauss et al., 2022), and therefore lends itself well to comparative questions. However, a comprehensive understanding of social relationships necessitates data on both affiliative and agonistic social interactions. Both types of interactions play crucial but distinct roles in shaping the dynamics and complexity of social systems. In contrast to dominance interactions, affiliative interactions are typically more varied in terms of both the types of behaviours and methods used to measure them, making it more complex to build a database of truly comparable data (Ellis et al. 2019). One approach to overcoming this challenge is to collate data for taxa where the observed behaviours and recording protocols are relatively uniform. These databases can then be integrated into an archive of databases, such as the EcoEvo data source catalogue (https://ckan-ecoevo.d4science.org/, Culina et al., 2018), to ultimately create a comprehensive FAIR database of social behaviour.

Here, we introduce MacaqueNet (https://macaquenet.github.io/), a global grass-roots collaboration that provides a platform for big team social behavioural research, and the MacaqueNet database, a cross-species collaborative FAIR database, essential for synthetic comparative research (Borries et al., 2016; Coles et al., 2022). To the best of our knowledge, this is the first collaborative database on both affiliative and agonistic social behavioural data in any animal and can thus provide a functional example for future endeavours of this type.

## Methods and Results

MacaqueNet connects researchers studying the social behaviour of macaques in diverse settings. The aim of MacaqueNet is twofold: to encourage and facilitate collaboration between independent research teams and to provide a lasting central platform for data archiving, standardising, and accession. Here we describe the establishment of MacaqueNet, from starting a large-scale collective to the creation of a cross-species collaborative database and the implementation of data entry and retrieval protocols. First, however, we illustrate the exceptional suitability of macaques as a taxon to initiate the construction of a cross-species database of social behavioural data.

### Why the genus *Macaca* lends itself to the creation of a comparative database of social behaviour

With 25 currently recognised extant species, *Macaca* is the most widely geographically distributed non-human primate genus (Roos et al., 2019). Macaques are one of the most versatile and adaptable primates, exploiting very different environments, from the temperate, mountainous habitats of Morocco and Japan to the tropical forests of Southeast Asia (Thierry, 2007). Throughout this range they live in areas of varying anthropogenic impact, from urban cities to natural habitats coming under increasing tourist and agricultural pressure (Radhakrishna et al., 2012). This variation in habitats and climates is reflected in the species’ ecology, mating system and social structure (Cords, 2012). All macaques are frugivorous, but their diet also includes other plant materials, including leaves, seeds and flowers, as well as small vertebrate and invertebrate animals. They range from being arboreal to semiterrestrial, and encounter different levels of predation (Fleagle, 2013). Mating is polygynandrous and reproduction can be seasonal or year-round, with substantial variation in male reproductive skew (biased distribution of offspring sired towards a few, usually high-ranking, males) across species (Ostner et al., 2008).

Despite this variation, macaques have a rather conserved social organisation. They typically live in relatively large multi-male multi-female groups of approximately ten to a hundred individuals in the wild and up to several hundred in urban or provisioned populations (Cords, 2012). Females are philopatric and form the core of the group, clustered into multigenerational matrilines. Males typically disperse from their birth group around puberty to breed elsewhere and may even change groups several times in their life (De Moor et al., 2020). Both male and female macaques exhibit a wide range of social behaviours, many of which are shared across all species (Thierry et al., 2007). While social relationships tend to be kin biased and hierarchical to some extent, macaques show substantial variation in the patterns of affiliative and agonistic interactions. As such macaques show an extraordinary example of diversity within unity, the evolution and underpinnings of which still elude us (Balasubramaniam et al., 2018; Thierry, 2007).

Many species of macaques have been extensively studied in the wild or naturalistic semi-free ranging settings for several years, yielding valuable insights into their behaviour (*Macaca assamensis*, Ostner & Schülke, 2018; *Macaca fascicularis*, Van Noordwijk & Van Schaik, 1985; *Macaca fuscata*, Nakagawa et al., 2010; *Macaca leonina*, Albert et al., 2013; *Macaca maura*, Okamoto et al., 2000 & Riley et al., 2014; *Macaca mulatta*, Cooper et al., 2022; *Macaca nemestrina*, Ruppert et al., 2018; *Macaca nigra*, Duboscq & Micheletta, 2023; *Macaca radiata*, Sinha, 2005; *Macaca sinica*, Dittus, 1975; *Macaca sylvanus*, McFarland & Majolo, 2013; *Macaca thibetana*, Li et al., 2020). Japanese macaques (*Macaca fuscata*) were one of the first species for which researchers recognised and followed subjects individually, generating unprecedented levels of detail on their behaviour. The first published record on macaque behaviour dates back to 1956 (Kawamura, 1956). Since then a body of research has leveraged macaque behavioural data, including comparative research across macaque species (Balasubramaniam et al., 2012; Balasubramaniam et al., 2020; De Waal & Luttrell, 1989; Sueur et al., 2011; Thierry, 2021), to greatly contribute to answering fundamental questions on the evolution, selective pressures and adaptive functions of social behaviour (Balasubramaniam et al., 2018; Brent et al., 2017; De Moor et al., 2020; Ellis et al., 2019; McFarland & Majolo, 2013; Micheletta et al., 2012; Riley et al., 2014; Schülke et al., 2010; Singh et al., 2010; Testard et al., 2021; Van Noordwijk & van Schaik, 1985; Young et al., 2014). Research on macaque behaviour and socio-ecology has also helped to ignite interest for subsequent investigations into other species. Research on kin biases (Brent et al., 2017; De Moor et al., 2020; Widdig et al., 2016; Yamada, 1963) and patterns of dominance and agonism (Balasubramaniam et al., 2012; Bernstein & Sharpe, 1966; Simons et al., 2022) are good examples of this. Building on these strong foundations, we believe that MacaqueNet can further promote collaborative and comparative research on social behaviour.

### Creating MacaqueNet

MacaqueNet originated in 2017 as a small-scale collaboration proposal for one specific study. From there, MacaqueNet developed and expanded to become a global grass-roots network of macaque researchers, with further potential collaborators being identified through word of mouth and a literature search on Google Scholar. We aimed to mitigate the inherent bias towards researchers from Europe and the US, and to facilitate the involvement of scientists from macaque-range nations and historically underrepresented research communities. An initial email was sent to all first and last authors of papers that contained or pointed to the existence of social behavioural data on macaques, asking if they would be willing to contribute those data to a collaborative database. Of the 193 researchers contacted, 68 did not reply or declined, 28 had no suitable data and 97 agreed to participate.

### The MacaqueNet Relational Database

We brought together social behavioural data from independent research teams into a standardised cross-species database. The MacaqueNet database currently contains social interaction data from 1981 to the present for 14 of the 25 recognised species of macaques (Fig 1). At the time of publication, the data have been collected on 21 wild, 22 captive, and 18 free-ranging populations (some of which have been observed for several study periods) at a total of 61 field sites, zoos, and research centres (Fig 2), resulting in social behavioural data on 3972 individual macaques (Fig 1). The full extent of the data available can be explored using the search tool on the MacaqueNet website: https://macaquenet.github.io/database/.

**Fig 1:**
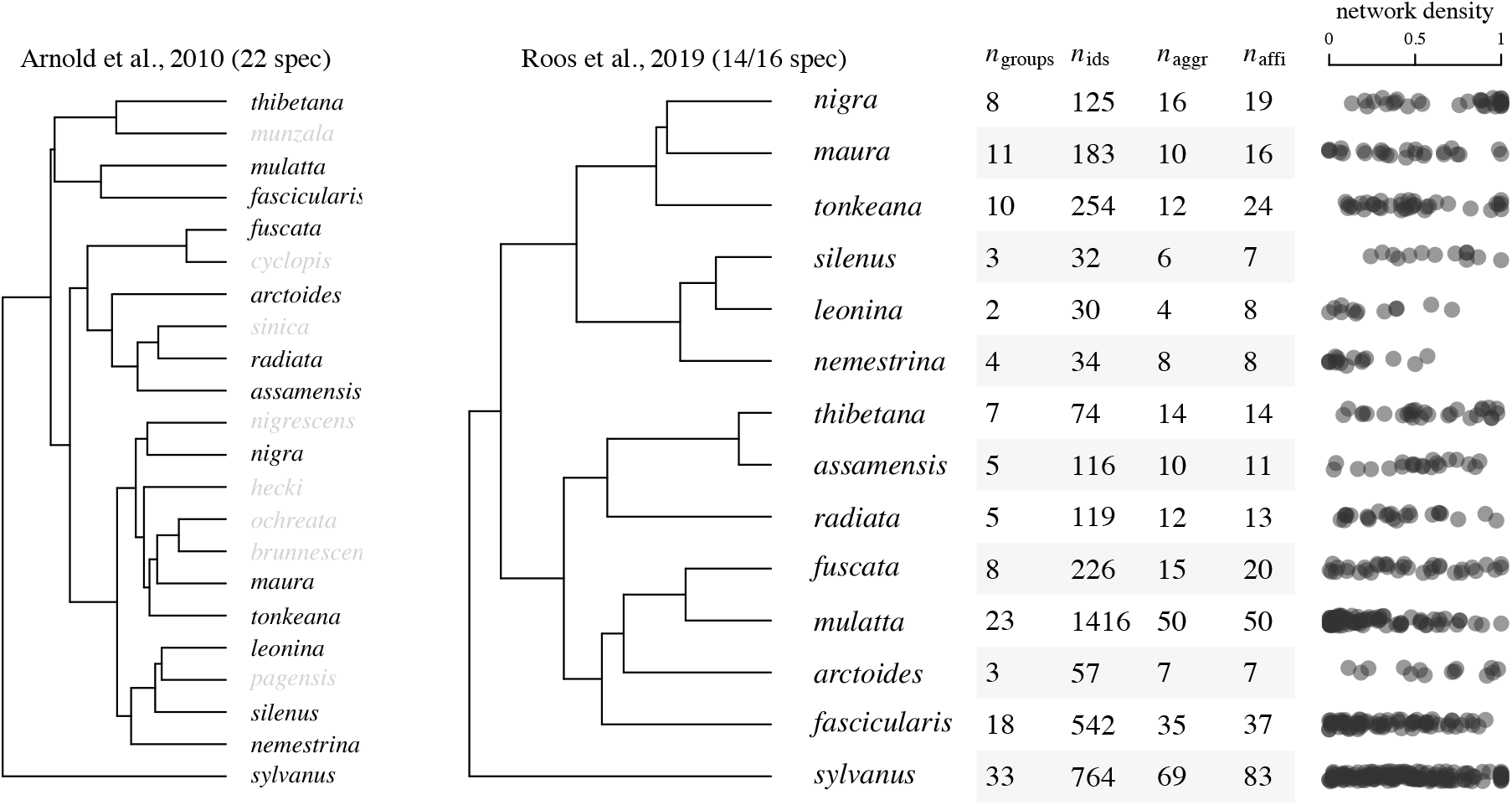
Summary of database content at the time of publication. The MacaqueNet database currently contains data for 14 out of 25 recognized species. The tree on the left shows the coverage of our species sample relative to the phylogeny with the highest species coverage (Arnold et al 2010). This tree does not include relatively newly described species: *Macaca selai, Macaca leucogenys* and *Macaca siberu*. On the right we show a more recent phylogeny pruned to our species sample (Roos et al 2019). The table contains the number of group-periods (i.e. a given study period for a given group), individuals and socio-metric matrices for aggression and affiliative behaviours for each species. As some groups and the individuals within them have been observed over multiple periods, the total count does not necessarily represent the number of unique groups or individuals. The dot plot on the right illustrates network densities (the proportion of dyads that were observed interacting at least once). Each dot represents the density for one (affiliative or agonistic) socio-metric matrix.

**Fig 2:**
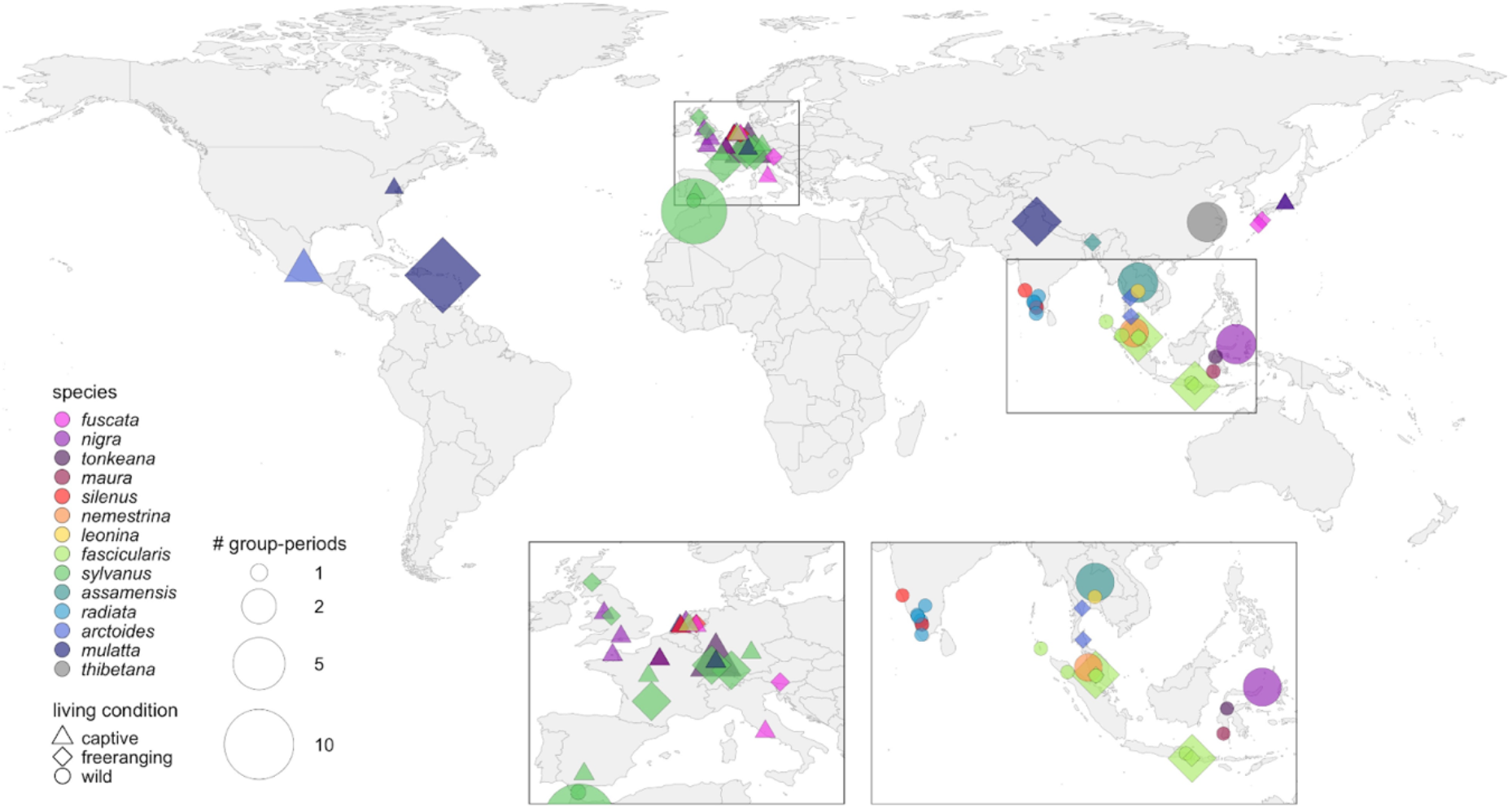
Geographical distribution of research sites in which the data currently present in the MacaqueNet database have been collected. Populations in America and Europe (with the exception of Gibraltar) have been introduced. Living conditions are classified as wild, free-ranging or captive (see Glossary in the supplementary material for definitions). Group-periods represent the total number of study periods for all groups at a given research site.

The core of the database consists of sociometric matrices representing the two primary axes of animal social structure: dyadic affiliative or agonistic behaviour between individuals, aggregated by study period (Fig 3A). The behavioural data comprise three categories of affiliative behaviour - grooming, spending time in body contact, and spending time in close spatial proximity - and two categories of agonistic behaviour - contact and non-contact aggression. These data represent the most common behavioural interactions expressed in macaques (Thierry et al., 2007), forming the backbone of macaque affiliative and dominance relationships, and are therefore collected by most researchers studying macaques. Data have been collected observing one focal individual or (part of a) group at a time, using continuous recording or discrete sampling of dyadic behaviours, recorded as counts or durations (see Glossary in the supplementary material for definitions). Each set of sociometric matrices for a particular group-period (i.e. a given study period for a given group) is accompanied by a subject data file, which contains individual attributes on sex, age, and observation effort (Fig 3B).

**Fig 3.**
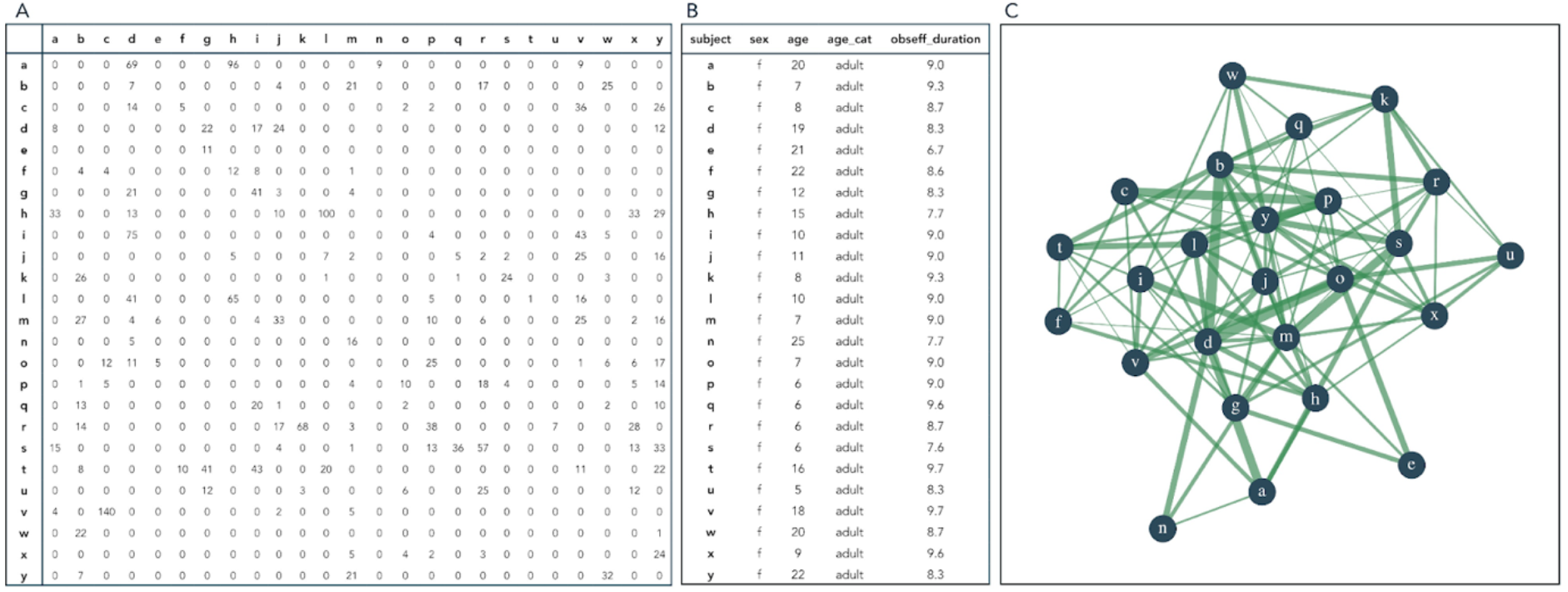
Example of the data of one dataset as stored in the MacaqueNet database. A) A sociometric matrix documenting pairwise interactions in a given group-period. For directed matrices, the actors - the individuals who initiate the behaviour - are listed in the rows, while the receivers - the individuals towards whom the behaviour is directed - are listed in the columns. The matrix entries represent either the total number of times (counts, as depicted here) or the total duration (in seconds) for which an individual in the row performed the behaviour towards the individual in the column for a given study period. B) A subject table listing all individuals observed in a given group-period, along with individual attributes: sex, age, and observation effort, here duration of observation in hours. C) An illustration of the network representing the data in the sociometric matrix. Each blue circle represents a subject, green lines between circles represent dyadic affiliation strength, here quantified as the dyadic rate of grooming.

In addition to the behavioural data, the database includes rich metadata on methodology, study populations and research teams. The data and metadata are organised as a relational database (see Fig S1 in the supplementary material for an overview of the relational database tables). All tables are linked by unique identifiers (the ‘-id’ columns in each table). Each row in the Behaviour table links to a sociometric matrix with all instances of a specific behaviour category for one group-period (e.g. counts of dyadic grooming events in a group of Barbary macaques (*Macaca sylvanus*) from La Montagne des Singes in 2017, Fig 3A). All variables are defined in the Glossary in the supplementary material.

To ensure that the data and metadata are comparable across studies, we set up a standardisation pipeline through which all contributed data (‘primary data’) is run before it enters the database. Fig S2 in the supplementary material provides an overview of the workflow from primary data submission, over the data checks and standardisation pipeline to data requests. Both the primary data and the standardised database are currently stored and managed in a GitHub repository (https://github.com/MacaqueNet). The code for the data standardisation pipeline and the metadata on the available datasets, along with resources such as the Glossary, the workflow and the MacaqueNet Terms of Use, are openly accessible in the repository.

### MacaqueNet is both a big-team science resource and a community

MacaqueNet makes macaque data FAIR (Findable, Accessible, Interoperable and Reusable, Wilkinson et al., 2016), in four main ways. First, by identifying, accessing, and bringing together data from multiple sources into one centralised place (https://macaquenet.github.io/database/). Second, by developing a common vocabulary for the data and metadata in the MacaqueNet database, with openly accessible definitions (see Glossary in the supplementary material, Borries et al., 2016). Third, by standardising data across studies and research sites together with the data contributors who understand the intricacies and subtleties of the data (Schneider et al., 2019). Fourth, by providing transparent and openly available annotation of how primary data were processed, so that the original data can always be recovered and processed in different ways if future researchers want to make other decisions (Schneider et al., 2019). As a result, MacaqueNet is a meticulously curated database with data readily available for future comparative studies (Borries et al., 2016; Gomes et al., 2022).

MacaqueNet extends beyond mere data sharing. By linking researchers across different research labs and institutions, MacaqueNet facilitates a global exchange of ideas and fosters the development of new research projects. To provide an opportunity for all the current data contributors to meet, discuss their research and learn about ongoing projects, we organised a virtual global symposium ‘Weaving the MacaqueNet’ in November 2021. During the symposium, future directions for MacaqueNet were discussed and agreed on. As a first step in the establishment of an infrastructure that permits researchers to communicate and coordinate their research, we set up a website (https://macaquenet.github.io), a mailing list and a group messaging channel on Slack. The second ‘Weaving the MacaqueNet’ symposium was held in August 2023 at the Joint Conference of the International Primatological Society and Malaysian Primatological Society, where the MacaqueNet community got the opportunity to meet in person. At that meeting, we discussed the expansion of MacaqueNet to include more types of data, thereby increasing MacaqueNet’s ability to address a range of questions on the biology of social behaviour. To promote transparency, we also agreed to make all meetings associated with MacaqueNet open to all MacaqueNet members, in addition to disseminating a quarterly newsletter outlining new projects and discussion points.

At the time of publication, four collaborative projects are using MacaqueNet data: 1) a project investigating the links between social diversity and social complexity; 2) a project exploring the socio-ecological drivers of variation in social relationships; 3) a project testing the impact of hierarchy steepness on rank-related benefits and 4) a project looking into predictors of intersexual dominance. New projects can be easily proposed, and data requested with a brief project proposal via the website (https://macaquenet.github.io/database/). The proposal is forwarded to the data contributors, who assess whether they can provide their data for the proposed project and determine their usage terms (see MacaqueNet Terms of Use in the supplement). New types of data to be added to the database can also be suggested on the website. These ideas and their potential implementation are discussed at each MacaqueNet symposium.

### The future potential of MacaqueNet

MacaqueNet’s potential to advance our understanding of social relationships extends beyond its current capability to explore their ultimate function and evolution. Macaques are one of the very few taxa that have been studied extensively both in the field and in the lab, making them an exceptionally well-suited comparative system for integrative studies. Macaques are the most used primate model in neurobiological and socio-developmental research (Capitanio & Emborg, 2008; Cooper et al., 2022), and the rhesus macaque (*Macaca mulatta*) genome – one of the most complete primate references (Warren et al., 2020) – is a crucial resource for new developments in the genetic causes and consequences of social behaviour (Chiou et al., 2020; Simons et al., 2022). To expand the scope of the MacaqueNet database, efforts are underway to include additional data types such as life-history, morphological, endocrinological, and genetic/genomic data. The MacaqueNet community is enthusiastic about sharing these data, which will enable new developments in the study of social relationships, their functions, and their underlying mechanisms.

We envision that MacaqueNet will have significant potential to shed light on four main areas of research on social evolution. First, the genus *Macaca* exhibits pronounced variation in ecology and social structure, but stable social organisation across species (Thierry, 2007). This makes it possible to quantify between- and within-species variation in social structure (Balasubramaniam et al., 2012; Neumann & Fischer, 2023) and explicitly test the social and ecological drivers of this variation. Important insights can be gained from combining both within- and between-species comparisons (Schradin, 2013; Sinha, 2005). Populations of the same species tend to differ in fewer characteristics, reducing the effects of confounding variables, while comparisons between species can reveal broader evolutionary patterns and processes (Rubenstein & Abbot, 2017).

Second, research from the lab can inform field research and vice versa (Simons et al., 2022; Testard et al., 2022), through a closer integration of neurological, physiological, genomic and behavioural research, to better capture the true complexity of the mechanistic underpinnings of social relationships in naturalistic settings. Third, a substantial amount of life-time life history data has been accumulated from multiple long-term field sites. This opens up the possibility to comparatively explore the impact of sociality on life-history, and to examine how individuals adapt their social behaviour in response to changing environments or needs throughout their lifespan (Siracusa, Negron-Del Valle, et al., 2022).

Finally, social behavioural data on macaques facing anthropogenic challenges, such as urbanisation (Balasubramaniam et al., 2020; Kaburu et al., 2019; Morrow et al., 2019), intensive farming (Holzner et al., 2019) and natural disasters (Testard et al., 2021) offers the unique opportunity to explore the impact of such challenges on individual health, aging, and fitness outcomes as well as group-level consequences such as changes in social structure and the increased risk of zoonotic disease spread. This research is essential for advancing population conservation efforts (Estrada et al., 2017) and transitioning human-wildlife interactions from conflict to coexistence (Radhakrishna et al., 2012).

## Discussion

In this paper, we advocate for the collaborative building of FAIR cross-species comparative databases as a critical tool in the study of the biological drivers of social relationships. Understanding the substantial variation in the social relationships animals form requires comparing taxa with distinct life histories, ecology, and interlinked evolutionary histories, all of which may be key drivers of social variation (Kappeler et al., 2019; Lukas & Clutton-Brock, 2018). Yet, despite decades worth of social behavioural data (Sheldon et al., 2022), substantial challenges have impeded systematic comparative research, one of which is the creation of truly comparable databases (Borries et al., 2016). We propose that large-scale collaborative efforts, where data and expertise are shared among researchers, can provide a solution to this issue and help facilitate reliable cross-species comparisons. To inspire and facilitate more collaborative efforts in the biology of social behaviour, we share our experience in developing MacaqueNet and its cross-species database.

Building on the foundations of previous big-team projects that have been instrumental in shaping MacaqueNet (including but not restricted to SPI-Birds: Culina et al., 2021 and ManyPrimates: ManyPrimates et al., 2019), we highlight three substantial advantages for the scientific community to join forces and pool their data into FAIR databases. First, by creating an enduring infrastructure for collaboration, big-team science efforts represent a crucial interface linking existing data and data users (Culina et al., 2021). They ensure that valuable data, which might have otherwise remained largely unused, are easier to discover, repurpose, and synthesise (Gonzalez & Peres-Neto, 2015; Purgar et al., 2022; Wallis et al., 2013) and help pinpoint areas that would benefit most from additional data collection.

Second, much of the existing social behaviour data are the result of years of effort and funding acquisition to set up the necessary logistics and to collect data on often wild and endangered animals. Big-team science can set up guidelines and regulations to ensure that sharing is done in such a way that it considers individual researchers’ interests, and their ability to acquire funding to keep collecting new data. Transparent but controlled accessibility, where data contributors retain full ownership of their data, best balances benefits to the scientific community while protecting data contributors, as proven successful in other big-team science endeavours (Culina et al., 2021; Strier et al., 2010). Creating a community rather than just a database also offers data contributors networking opportunities and access to others’ data, fostering collaborative research possibilities.

Third, social behavioural data are particularly complex, with many researcher degrees of freedom involved in how the data are defined and what methods of observation are used (Borries et al., 2016). MacaqueNet demonstrates the benefits of starting small, with a single taxon that exhibits similar social organisation and behaviour across species. This approach allowed us to focus on core issues, such as data comparability and standardisation, and identify potential solutions to address those issues. For instance, MacaqueNet has spurred the creation of BISoN, a novel Bayesian framework for inference of social networks (Hart et al. 2023). Two other projects have arisen to address challenges in comparative social network analysis that were long recognized but for which MacaqueNet has provided well-defined questions and the data necessary to address them. Our experience also emphasised the importance of standardising data through direct interaction with data contributors, thus ensuring clarity around the intricacies of the data and averting inadvertent misinterpretations. These developments, along with the established infrastructure, such as the guidelines, workflow and code for dealing with data sharing, can pave the way for databases of larger taxonomic scope and including more types of data. This will be key to get a comprehensive understanding of the development, regulating mechanisms and life-time fitness consequences of social relationships framed in the context of relevant socio-ecological pressures and phylogenetic history.

The rise of collaborative databases in a variety of fields is a testament to the enthusiasm of researchers to join forces and share data and resources towards comparative and interdisciplinary studies, an emerging and promising way of doing science (Coles et al., 2022). Big-team endeavours like MacaqueNet facilitate the exploration of innovative and comparative questions no single research lab could address individually and make testing the reproducibility of research findings across contexts possible. While the creation of databases on social behavioural data comes with substantial challenges, we truly believe this is the way forward to harness the incredible comparative potential of the wealth of existing social behavioural data. We hope that MacaqueNet, along with similar endeavours, can catalyse large-scale collaborative research into the biology of social behaviour.

## Supporting information

Glossary

MacaqueNet Terms of Use

## Supplementary material

Fig S1: Set-up of the relational MacaqueNet database. All variables are defined in the Glossary. Arrows indicate how each table is linked to at least one other table in the database, through unique identifiers (the ‘-id’ or ‘running_number’ columns in each table). Variables can be of the data type ‘text’, ‘num’ (numerical), ‘int’ (integer), ‘cat’ (categorical), ‘date’ or ‘bit’ (binary digit, 0/1). Part of the behaviour and subject tables are only relevant for the data import pipeline. The ‘sociometric_matrix’ table refers to sociometric matrices with all instances of a specific behaviour category for one group-period rather than an actual table.

Fig S2: Overview of the workflow from primary data submission, over the data checks and standardisation pipeline to data requests. All contributed data contain several sociometric matrices along with corresponding subject data and metadata for at least one group-period for a specific species. The appropriate subset of the database to be used will vary depending on the planned projects by the data users. For instance, let’s consider three datasets; data 1 has information on grooming, proximity and non-contact aggression in a group of rhesus macaques, data 2 has information on grooming and body contact in another group of rhesus macaques, and data 3 has data on grooming, body contact, and contact aggression in a group of Barbary macaques. If data user 1 is planning a project on rhesus macaques’ affiliation, they would only use data 1 and data 2. On the other hand, if data user 2 is planning a project on body contact across different species, they would only use data 2 and data 3.

## Notes

https://macaquenet.github.io/

https://github.com/MacaqueNet/

### Competing Interest Statement

The authors have declared no competing interest.

