## Supplementary material for "MacaqueNet: big-team research into the biological drivers of social relationships": Glossary

### MacaqueNet Glossary

Within the MacaqueNet database, a dataset is defined as an interaction matrix with the corresponding subject data, associated metadata on data collection and metadata specific to the behaviour in the matrix for a given group-period. Each dataset consists of several unique dataframes. Each dataset consists of several unique dataframes. Each dataframe is either a socio-metric matrix of a specific social behaviour or the subject data for a given group-period.

#### research site

***research_site* -** the name of the research centre, zoo, animal park or field site at which each dataset was collected and by which it is known in the literature.

***site_id* -** a unique number randomly assigned to each unique research site where data were collected.

***country_site* -** the country in which each research site is located.

***latitude_site* -** the latitudinal coordinates for each unique research site, given in degrees and rounded to the nearest integer.

***longitude_site* -** the longitudinal coordinates for each unique research site, given in degrees and rounded to the nearest integer.

#### species

***species_common* -** the common name for each unique macaque species, as specified by the IUCN (International Union for Conservation of Nature).

***species_scientific* -** the scientific name for each unique macaque species. This is the last part of the latin name (specific epithet), not including the genus name.

***species_id* -** a unique number randomly assigned to each unique macaque species.

#### population

***pop_id* -** a unique number randomly assigned for each unique population, defined as a unique combination of species and research site.

#### group

***group_name* -** the name of the group within a population on which each dataset was collected and by which it is known in the literature. Group_name is consistently used for the same social group, even if different names were given by different teams or in different data collection periods. If only one group exists within a population, Group_name is 'onegrouponly'.

***other_names*** - alternative names used in the literature for a given group, separated by a semi colon (;). If there are no other names for a unique group, the other_names column remains blank.

***group_id* -** a unique number randomly assigned to each unique social group.

***range_management*** - the extent to which a group's range is managed. Can be one of three options: "wild" (animals can move freely between different groups or populations, without range restrictions), "free-ranging" (animals can move freely between different groups or populations, but are confined to a specific area intentionally by humans) or "captive" (animals have restricted ranges and cannot move freely or disperse between different groups or populations).

***medical_intervention* -** whether a group receives medical interventions in the form of reproductive control and/or treatment of injury or disease as part of the population's management plan. Can be one of two options: "1" (routine medical interventions) or "0" (no routine medical interventions).

***food_provisioned* -** whether humans routinely and intentionally provide a group with food. Can be one of two options: "1" (food provisioned) or "0" (not food provisioned).

***human_contact* -** whether a group is in regular contact with non-researcher humans, such as groups whose ranges overlap with human settlements or groups regularly visited by tourists. Can be one of two options: "1" (regular contact with non-researcher humans) or "0" (no regular contact with non-researcher humans).

#### group_period

***group_period_id* -** a unique number randomly assigned to each unique group-period, defined as a unique combination of social group and study period.

***start_year -*** the year in which data collection commenced for each unique dataset, denoted in 4-digit format.

***start_month -*** the month in which data collection commenced for each unique dataset, denoted as the first 3 letters of the month name.

***end_year -*** the year in which data collection concluded for each unique dataset, denoted in 4-digit format.

***end_month -*** the month in which data collection concluded for each unique period, denoted as the first 3 letters of the month name.

***group_size -*** the total number of individuals in each social group during the period of data collection. This value includes all group members: (sub)adults, juveniles and infants. If group size varies across the period of data collection, the largest group size is noted. If the exact group size is not known, the group_size column remains blank.

***adult_males -*** the total number of adult males in the group at the time data were collected. This value includes males who were or became adults during the study period. If the number of adult males in the group varies across the period of data collection, the largest number of adult males is noted.

***adult_females -*** the total number of adult females in the group at the time data were collected. This value includes females who were or became adults during the study period. If the number of adult females in the group varies across the period of data collection, the largest number of adult females is noted.

#### behaviour

***running_number -*** a unique number randomly assigned to each unique dataframe for each unique dataset.

***folder -*** the name of the folder in which the original data (as provided by the researcher who contributed the data, before going through the standardisation pipeline) is stored within the primary data repository, denoted as the species' scientific name and the last name of the lead researcher who contributed the data to the database.

***file -*** the name of the file containing the original data, as provided by the researcher who contributed the data to the database.

***sep -*** the format of each original dataframe. Can be one of four options: “tab” (for dataframes shared as tab separated files such as .txt files), “comma” (for dataframe shared as comma separated files, such as .csv files) “excel” (for dataframes shared as Excel sheets) or “manual” (for datafiles that had to be entered manually).

***sheet -*** for datasets shared as Excel files with multiple sheets. Denotes the name of the sheet containing each dataframe, as provided by the researcher who contributed the data to the database. If data were not shared as Excel files, the sheet column remains blank.

***sheetrange -*** for primary datasets that contain multiple dataframes (e.g. behavioural matrices) on a single sheet. Denotes the cell range containing each dataframe. If data were not shared as Excel files, the Sheet column remains blank.

***cat_global -*** the broad data category of the dataframe. Can be one of three options: "affi" (affiliative) or "aggr" (aggressive) for behavioural data or "subjectdata" for subject data.

***cat_sub -*** the specific data subcategory. Can be one of six options: If cat_global is "aggr" (aggression), cat_sub can be "conagg" (aggression that involves contact), "nonconagg" (aggression that does not involve contact) or "allagg" (if aggression type is not specified or if both contact and non-contact aggression are combined into one matrix). If cat_global is "affi" (affiliation), cat_sub can be "bodycon" (body contact), "groom" (grooming) or "prox" (time spent in proximity).

***cat_detail -*** provides more detail regarding the cat_sub column: If cat_sub is "conagg", cat_detail will be e.g. "hit", "bite". If cat_sub is "nonconagg", cat_detail will be e.g. "chase", "subm" (submissive), "displ" (displacement) or "suppl" (supplants).

***focal_data -*** whether data were collected via focal animal sampling or group sampling. Can be one of two options: “1” (focal animal sampling) or “0” (group sampling).

***sampling_method -*** the sampling method used to record the behaviour. Can be one of two options: "discrete" (recording at regular time intervals whether the behaviour of interest is occurring, sometimes also called ‘time sampling’, ‘instantaneous sampling’ or ‘scan sampling’) or "continuous" (recording all instances of a behaviour continuously, sometimes also called ‘all occurrence sampling’).

***data_type -*** the format of the data in each dataset. Can be one of two options: “count” (the number of occurrences of a behaviour, if the behaviour was recorded continuously, or the number of discrete samples in which the behaviour was occurring, if the behaviour was recorded discretely) or “duration” (the length of time a behaviour occurred for).

***interval -*** for data collected using discrete sampling. Denotes the interval between discrete samples, in seconds.  If data were collected continuously, the interval column remains blank.

***obseff -*** the format of the observation effort Can be one of two options: “Obseff_samples” (observation effort is expressed as a number of time samples) or “Obseff_duration” (observation effort is expressed as a duration).

***corrected_obseff -*** whether the data shared by the data contributor are corrected for observation effort.

***data_symmetric -*** whether a sociometric matrix is symmetrical or not. Can be one of two options: “0” (the matrix contains directed behaviour with a clear actor and receiver, and is not symmetrical), or “1” (the matrix contains undirected behaviour, and is symmetrical).

#### subject_data

***nr_ama_subjects -*** the number of adult male subjects. This is the total number of adult males for which behaviour was recorded in the dataset for a specific period.

***nr_afe_subjects -*** the number of adult female subjects. This is the total number of adult females for which behaviour was recorded in the dataset for a specific period.

***all_adults_obs -*** whether or not all adults who were present in the group during the period of data collection were observed. Can be one of two options: “1” (yes, all adults in the group were observed) or “0” (no, not all adults in the group were observed).

#### subjects

***subject -*** the identification name of each subject from which data were collected for each dataset, as specified by the researcher who contributed data to the database .

***sex -*** the sex of each subject. Can be one of two options: “male” or “female”.

***age -*** the age of each subject in years, rounded to the nearest integer. The age column may be left blank if a subject's exact age is not known, but age category is known.

***age_cat -*** whether the subject is an adult, subadult, juvenile or infant. Age categories are based on the age cutoffs assigned by the data contributor and therefore, age ranges for each category might vary between datasets collected on the same group, the same species or at the same field site.

***obseff_samples -*** for data collected using discrete sampling only. Denotes the observation effort for each unique subject, expressed as a number of discrete samples.

***obseff_duration -*** for data collected using continuous sampling only. Denotes the observation effort for each unique subject, expressed as a duration in hours.

#### researcher

***lead_contact*** - the last name of the corresponding person who serves as the primary point of contact for each dataset.

***researcher_first -*** the first name of each researcher who contributed data to the database.

***researcher_last -*** the last name of each researcher who contributed data to the database.

***researcher_id -*** a unique number randomly assigned to each unique researcher who contributed data to the database. Each member of a team who contributed to the creation of each unique dataset will have a unique Researcher_id.

***researcher_email -*** the current email address of each researcher who contributed data to the database.

#### affiliations

***institute*** - the name of the institution(s) that each researcher who contributed data to the database is currently affiliated with.

***institute_id*** - a unique number randomly assigned to each unique institute that each researcher who contributed data to the database is currently affiliated with.

***country_inst -*** the country in which each unique institution to which each data contributor is affiliated is located.

***latitude_inst -*** the latitudinal coordinates for each unique institution to which each data contributor is affiliated, given in degrees and rounded to the nearest integer.

***longitude_inst -*** the longitudinal coordinates for each unique institution to which each data contributor is affiliated, given in degrees and rounded to the nearest integer.

#### team

***team_id -*** a unique number randomly assigned to each unique team of researchers for each dataset. This is each unique combination of researchers who jointly contributed the same dataset to the database. A single researcher may be a member of multiple teams. The decision as to who is considered a member of the team for each dataset is determined by the data contributor.

#### research_project

***project_name -*** the full name of each unique project using the MacaqueNet database, as it is named on the MacaqueNet website.

***project_id -*** a unique number randomly assigned to each unique research project that is using the MacaqueNet database.

***project_contact_first -*** the first name of the person who serves as the primary point of contact for each research project.

***project_contact_last -*** the last name of the corresponding person who serves as the primary point of contact for each research project.

***project_contact_email -*** the email address of the corresponding person for each research project.

#### permissions

***metadata_access -*** the level of access to the metadata of each dataset, as selected by the data contributor. Can be one of two options: “open” (open access to the metadata) or “onrequest” (access to the metadata can be requested by a researcher wishing to utilise the data for their project).

***data_access -*** the level of access to the behaviour or subject data of each dataset, as selected by the data contributor. Can be one of two options: “open” (open access to the data) or “onrequest” (access to the data can be requested by a researcher wishing to utilise the data for their project).

***permission_id -*** denotes the access levels for each unique dataset. Can be one of four options, representing each unique combination of metadata_access and data_access: “1” (open access to both the metadata and the data), “2” (both metadata and data are only available on request), “3” (open access to metadata, but data only available on request).

#### project_data

***access_granted -*** whether the researcher who contributed data to the database has granted permission for their dataset(s) to be used in each specific research project. Can be one of 2 options: “1” (permission has been granted) or “0” (permission has not been granted).
