## Supplementary figures and images for "MacaqueNet: big-team research into the biological drivers of social relationships"

### Fig S1

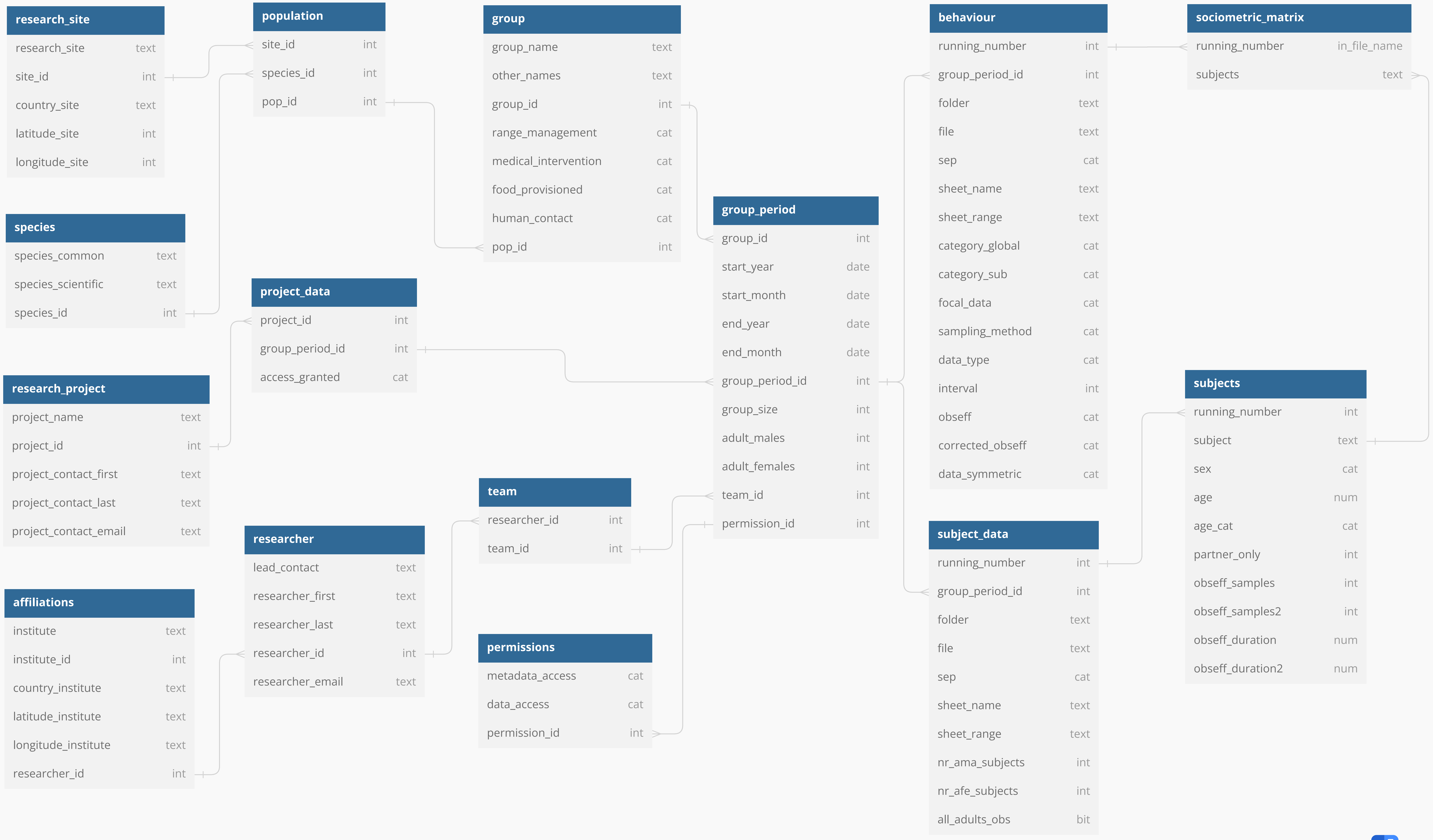


dbdiagram.io

### Fig S2

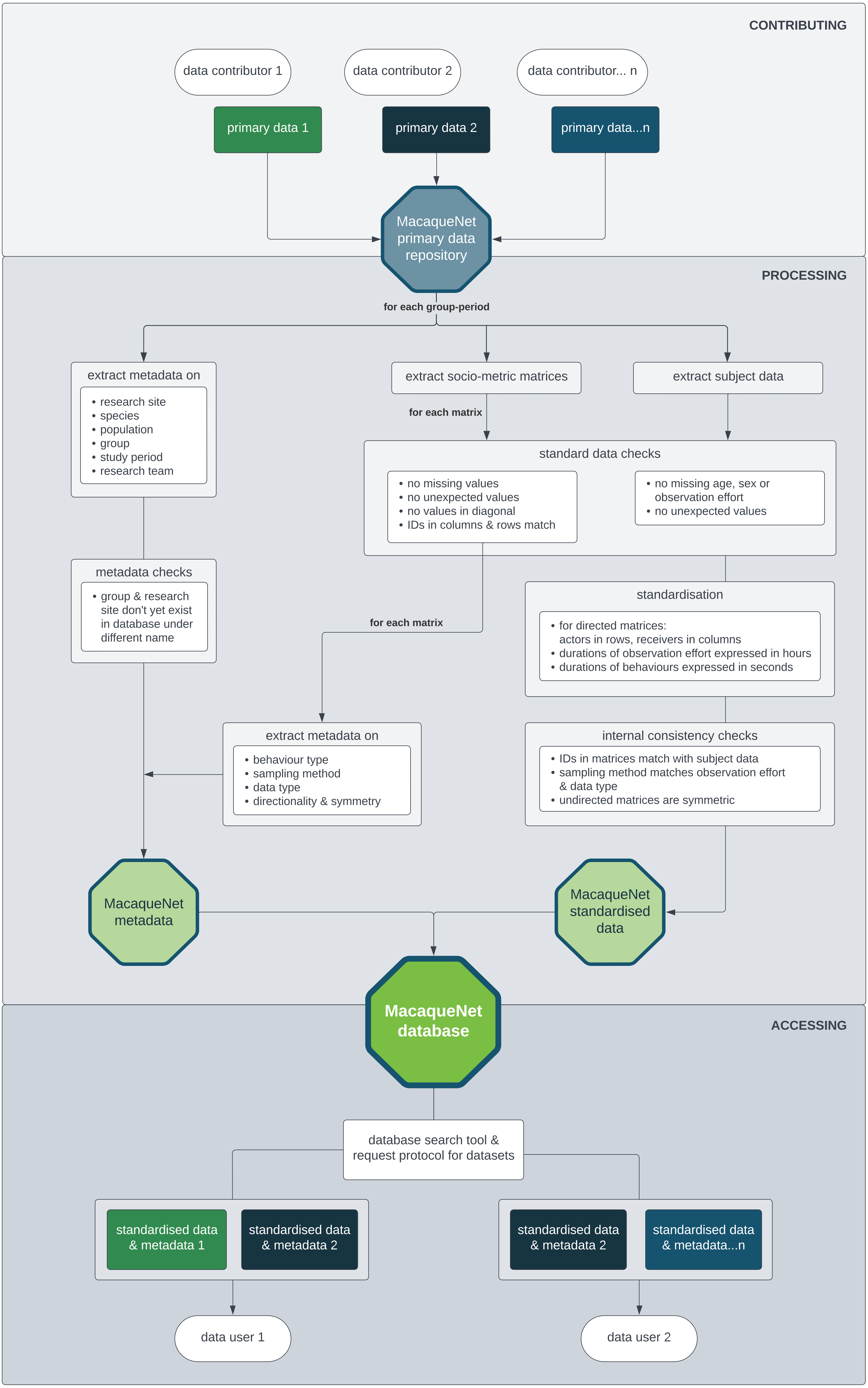
