## Supplementary material for "MacaqueNet: big-team research into the biological drivers of social relationships": MacaqueNet Terms of Use

### Terms of Use

Version 1.0.0

10 August 2023

Acknowledgements:
This terms of use document was adapted (with permission) from the SPI-Birds Network and Database. We would like to thank Antica Culina for her guidance in producing these terms of use.

1. INTRODUCTION

This document was created for MacaqueNet. MacaqueNet is a global grassroots network, whose mission is to encourage and facilitate collaboration between macaque researchers. MacaqueNet aims to strengthen and create new connections between macaque researchers, to facilitate data sharing and promote truly interdisciplinary and cross-species big-team science. Together, we can join forces to establish a culture where knowledge and resources are shared to let comparative macaque research reach its full potential.

This terms-of-use document provides a guide to contributing, accessing and using data stored in the MacaqueNet database. Each dataset is accompanied by metadata, which provides information including data contributors, population, study period, subject names as well as the level of access (open or accessible upon request) for both the metadata and data of each dataset, as selected by the data contributor. We encourage the use of the MacaqueNet database for academic, research, education, and other non-profit professional purposes. Access to any dataset stored in the database may be requested for a specific project. Users must read and agree to MacaqueNet's data use agreement below, and adhere to the principle of acknowledging the use of data and its attribution.

2. GLOSSARY

| TERM | DEFINITION |
|---|---|
| data contributor | The person, team or organisation that owns or has custody of a dataset, and contributed it to the MacaqueNet database. |
| lead_contact | The person who serves as the primary point of contact for each dataset. |
| data user | The person or team who is requesting access to use a dataset or datasets stored in the MacaqueNet database. |
| data_access | The level of access to the behaviour or subject data of each dataset, as selected by the data contributor. Can be one of two options: "open" (open access to the data) or "onrequest" (access to the data can be requested by a researcher wishing to utilise the data for their project). |
| usage terms | Specific terms for using a dataset, as prescribed by the data contributor, often in correspondence with a data user, specifying any additional conditions under which their data may be used. |

3. DATA USE REQUIREMENTS

Access to datasets stored in the MacaqueNet database may be granted to a data user, depending on the data_access level of the dataset and the outcome of the correspondence between the data user and data contributor for closed datasets.
The general MacaqueNet terms of use apply to all users downloading data that are not their own. By accepting this document and accessing data from the MacaqueNet database, the user agrees to the following:

1. To follow the terms defined in the data_access level, and the study's usage terms for each study for which data are accessed and utilised.

2. To obtain permission from the data contributor for the proposed use to accessing and utilising data. For datasets with the data_access level set to 'open', permission of the data contributor is not required for use. However, we strongly encourage data users to make a reasonable attempt to contact the lead_contact about their intended use, and to offer co-authorship or other credit as appropriate for publications or products resulting from the use of a dataset.

3. To use the data only after obtaining explicit permission granted specifically for the intended project. If the data is intended for new applications or analyses, or if the applications or analyses significantly differ from the originally proposed project, permission from the data contributor needs to be obtained again.

4. To follow MacaqueNet citation guidelines (Annex I) for citing the use of data from the MacaqueNet database. Citation for individual datasets is only required if specified in the usage terms, as agreed with each data contributor. However, we encourage following these guidelines or providing citation and acknowledgement as is feasible.

5. Not to hold MacaqueNet or the data contributor liable for errors in the data. We do not guarantee the completeness or accuracy of the datasets stored in the MacaqueNet database.

6. To assess the suitability of each dataset for its intended use. Users are responsible for determining the suitability of each dataset they request access to. Allowing the download of these data does not suggest that MacaqueNet or the data contributor approve of the proposed use or support the results of your analysis. If you find potential errors or incomplete information within a study, please contact the lead_contact and/or.

4. DISCLAIMER

By using these data, the data user agrees to follow the guidelines outlined in this agreement. While substantial efforts are made to ensure the accuracy of data and documentation, complete accuracy of datasets cannot be guaranteed. MacaqueNet shall not be liable for damages resulting from any use or misinterpretation of datasets. The data user may be held responsible for any misuse that is caused or encouraged by the data user's failure to abide by the terms of this agreement. Data users should be aware that we periodically update datasets. Thank you for your understanding and cooperation.

#### ANNEX I

The following guidelines and examples are provided for data users, data contributors and publishers to ensure appropriate citation of data hosted at the MacaqueNet Database. Because data contributors retain custody of their data in the MacaqueNet database, references to "MacaqueNet data" or "data accessed through MacaqueNet" are insufficient. Proper citation or other attribution will depend on the data contributor's terms of use (as agreed with each data contributor for a specific study).

1. Cite MacaqueNet
Every paper using data from MacaqueNet should cite the MacaqueNet database itself, using this citation: De Moor et al. 2023 [add paper pre-print].
We also suggest including MacaqueNet as an author, associating that name with a large contributor list (as done in the MacaqueNet paper, click here for the full list of affiliations). This approach acknowledges both MacaqueNet and data contributors without excessively extending the author list.

2. Acknowledge MacaqueNet funders
Every paper using data accessed via MacaqueNet should acknowledge MacaqueNet funders: a European Research Council Consolidator grant (FriendOrigins - 864461).

3. Recognition for data contributors
The data contributor might request that you cite related papers or research projects, and/or acknowledge funders, field stations, field teams, research councils, national parks, etc. This will be agreed upon between the data user and the data contributor in the terms of use for each dataset for a specific proposed project.

4. Cite datasets hosted at MacaqueNet
To refer to each dataset hosted by MacaqueNet, it is best to include the following metadata: the population and the group names, the study period, the behaviour, the "MacaqueNet ID", the data version and the date accessed. For example:

"The data used in this study were accessed from MacaqueNet (population: Cayo Santiago, group: F, study period: July-November 2012, behaviour: grooming, MacaqueNet ID: 1003, data version: 1) on August 8th 2023."

This reference format can also be used for data availability statements, as it follows the recommendations by Springer-Nature and a research data policy framework drafted by the Research Data Alliance (Hrynaszkiewicz et al. 2020).

5. Cite large numbers of datasets/publications

In collaborations involving many studies in MacaqueNet, such as meta-analyses or comparative analyses, the content format or journal might limit the number of references and make it impossible to cite all participating studies and existing papers as described above. Examples of ways to include a longer list of citations or alternative acknowledgements:

(a) Some journals allow a separate reference list for the methods that might allow a full listing of citations for participating studies.

(b) Web-based tools or products should include a page of acknowledgements or source information that lists citations as suggested above, or otherwise reference participating studies and data contributors.

(c) Including a full list in the supplemental materials of a paper should be used only as a last resort. References listed here are typically not recognized by search engines and other automated methods of recognizing data citation and re-use.

We welcome discussion with publishers and others about how to better support attribution for these types of projects. Please contact.
